# Decoupling Central Metabolism from Catabolite Repression Enables Robust Cellulosic Sugar Co-consumption in *E. coli*

**DOI:** 10.64898/2025.12.27.696699

**Authors:** Utsuki Yano, Payel Sarkar, Michael D. Lynch

## Abstract

Lignocellulosic biomass is the most abundant and sustainable carbon source for bioproduction, but its efficient utilization is hampered by the heterogeneous mixture of sugars released upon hydrolysis. Most industrial strains consume these mixed sugars sequentially due to strong regulation and cross-inhibition, leading to complex processes and reduced carbon efficiency. To address this, we leverage a novel central metabolism that decouples central carbon flux from native regulatory feedback by employing a Gluconate-Bypass of glycolysis. We demonstrate that the Gluconate-Bypass effectively alleviates feedback regulation in *E. coli,* enabling co-consumption of glucose and xylose. Further strain engineering leads to the first robust co-utilization of four major lignocellulosic sugars: glucose, xylose, arabinose, and galactose. By decoupling central carbon flux from native regulatory feedback, this architecture provides a feedstock-agnostic platform that maintains high and robust consumption regardless of extreme fluctuations in sugar composition.

## Introduction

Metabolic engineering is key to converting renewable carbon sources into value-added products.^1^ Among these, lignocellulosic biomass is the most abundant and sustainable raw material, as it does not compete with global food supplies.^2,3,4,5^ Hydrolysis of lignocellulose yields a complex mixture of simple sugars, primarily glucose (from cellulose) and the pentoses including xylose, arabinose, and galactose (from hemicellulose).^6,7^ However, the commercial viability of lignocellulose-based biorefining is limited by the inefficient co-utilization of these mixed sugars and the large variability in commercial sugar composition. *Escherichia coli*, a key industrial host, exhibits strong carbon catabolite repression (CCR) driven by glucose uptake systems and glycolytic regulation. This global regulatory mechanism prioritizes glucose consumption and suppresses the simultaneous utilization of other sugars,^8–10^ leading to sequential sugar consumption that prolongs fermentation and reduces carbon efficiency. (Figure 1a).

**Figure 1.**
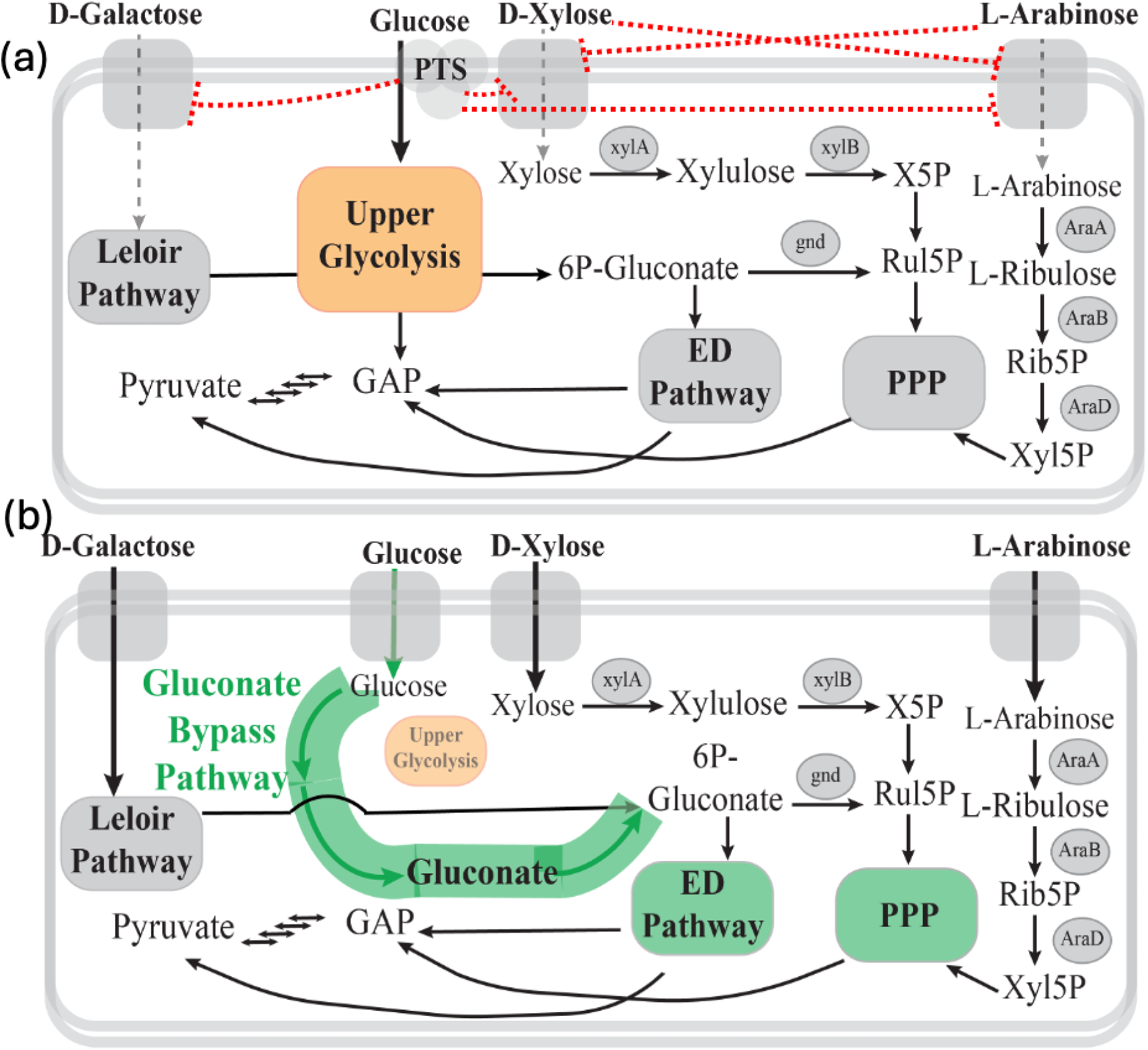
Comparison of sugar metabolism and regulatory networks in native *E. coli* and the engineered gluconate bypass strain. (a) Native *E. coli* central metabolism and carbon catabolite repression (CCR). The red dotted lines indicate inhibition of non-glucose sugar uptake and metabolism caused by PTS-dependent glucose transport, as well as additional cross-inhibitory interactions between sugar pathways. (b) The GBP architecture bypasses upper glycolysis by uptaking glucose through a PTS-independent system and converting it to gluconate, which enters the pentose phosphate (PPP) and Entner–Doudoroff (ED) pathways, placing all major sugars into shared metabolic routes. Because the GBP architecture is decoupled from glycolytic regulatory control, it enables robust co-utilization of glucose, xylose, arabinose, and galactose even under substantial variation in lignocellulosic hydrolysate composition.

Traditional strategies have failed because they often attempt to accommodate or mitigate existing regulatory constraints. This can leave strains vulnerable to the heterogeneity of sugar composition inherent in sustainable biomass feedstocks. While prior work has targeted key regulatory mechanisms to partially alleviate CCR, these often compromise growth or fail to achieve robust co-utilization of all lignocellulosic sugars simultaneously.^11–15^ Co-culture strategies, in which individual strains are engineered to consume specific sugars, can enable co-utilization but complicate process control and industrial scale-up.^16–20^ Introducing non-native pentose utilization pathways (e.g., Weimberg and Dahms pathways) offers simultaneous utilization potential, but these systems often lack robustness and are highly sensitive to variations in sugar composition.^21–23^ While achieving co-consumption of all lignocellulosic sugars is an important advancement, the ability to adapt to variations in sugar composition is equally critical. Lignocellulosic hydrolysate compositions vary widely depending on feedstock source and pretreatment method,^24,25^ making robustness, defined here as stable performance across diverse feedstock compositions, essential for industrial scale up. To our current knowledge there are no industrial strains capable of robustly co-consuming lignocellulosic sugars, and therefore the development of such strains remains a key unmet need.

To address the limitations of cross-inhibitory constraints, we leveraged a synthetic central metabolism, known as the gluconate-bypass (GBP). The GBP architecture strategically reroutes carbon flux to bypass glycolysis: glucose is transported via a PTS-independent system, oxidized to gluconate, and subsequently feeds into the Pentose Phosphate (PPP) and Entner–Doudoroff (ED) pathways (**Figure 1b**). Given our previous demonstration that this pathway circumvents glycolytic regulatory feedback driven by pyruvate accumulation,^26^ we hypothesized that this decoupled central metabolic network could similarly circumvent the regulatory constraints of CCR and enable simultaneous consumption of diverse sugars.

In this work, we demonstrate that the GBP metabolism alleviates the natural CCR in *E. coli.* We further report systematic engineering to resolve additional cross-inhibition among non-glucose sugars,^13,27,28^ resulting in a strain capable of fully robust co-utilization of glucose, xylose, arabinose, and galactose. To our knowledge, this is the first instance where glucose metabolism has been strategically bypassed to deregulate central carbon metabolism and achieve robust co-utilization of the diverse sugar components present in lignocellulosic hydrolysates. Importantly, we validate that the GBP architecture enables robust, balanced co-utilization of all major lignocellulosic sugars, even under substantial variation in feedstock composition.

## Results

### Glucose & Xylose Co-Utilization with Gluconate-Bypass Metabolism

The GBP metabolism bypasses upper glycolysis by converting glucose into gluconate-6-phosphate that feeds into the PPP and ED pathways (Figure 1b). As a result, we hypothesized that this re-wired central metabolic architecture could enable simultaneous utilization of glucose and xylose, the primary C5 sugar in lignocellulosic hydrolysates. To test this, we evaluated the strain engineered with GBP central metabolism, DLF_GBPC1, in minimal media containing glucose and xylose at a 2:1 ratio, mimicking their relative abundance in in several lignocellulosic hydrolysates.^29^ As shown in Figure 2a, DLF_GBPC1 co-consumed both sugars, whereas the control strains DLF_Z0025 (PTS-dependent, glycolysis-based glucose metabolism) and DLF_0286 (non-PTS-dependent, glycolysis-based metabolism) displayed the expected glucose-preferential consumption. (A schematic comparison of carbon metabolism in the two control strains is provided in Supplemental Figure S1.) We next quantified sugar uptake rates in DLF_GBPC1 (Figure 2b). DLF_GBPC1 consumed glucose and xylose in parallel, with glucose consumed approximately 1.4-fold faster than xylose (0.87 vs. 0.62 g L⁻¹ h⁻¹), demonstrating the suitability of this strain for lignocellulosic hydrolysates.

**Figure 2.**
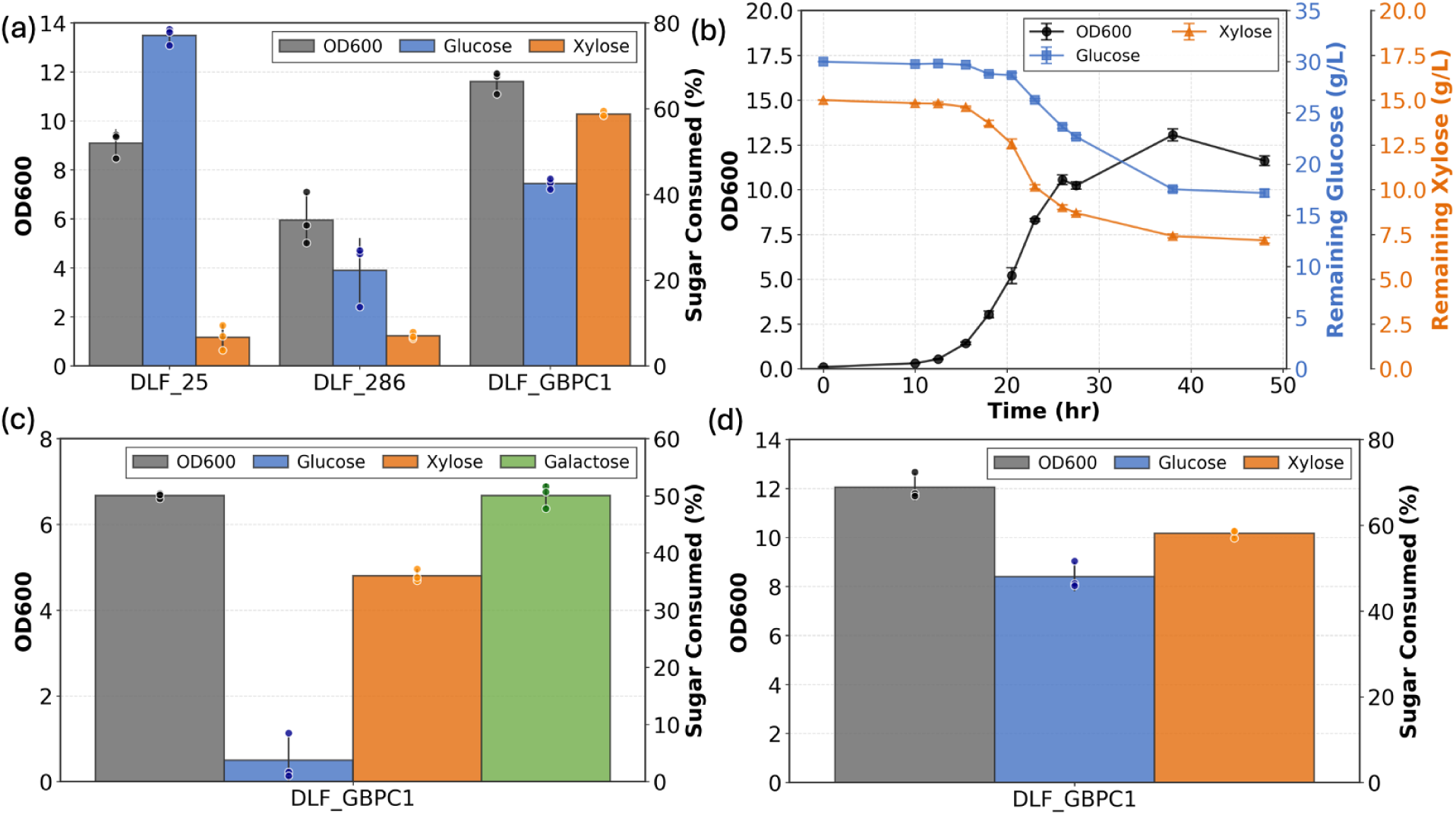
Co-utilization of glucose and xylose by DLF_GBPC1 strain. (a) Comparison of biomass (black), glucose consumption (blue), and xylose consumption (orange) in PTS-dependent (DLFZ_25), PTS-independent (DLF_286), and GBP strain expressing GalP (DLF_GBPC1) grown in media containing glucose and xylose at a 2:1 ratio (30 g L⁻¹ glucose, 15 g L⁻¹ xylose) (b) Time-course profiles of growth and glucose and xylose consumption for DLF_GBPC1 in glucose–xylose media. (c) Consumption of glucose (blue), xylose (orange), and galactose (green) by DLF_GBPC1 in GXAG media containing glucose, xylose, arabinose, and galactose at a ratio of 23.5:12.5:2:1 (27.03 g L⁻¹ glucose, 14.38 g L⁻¹ xylose, 2.31 g L⁻¹ arabinose, 1.15 g L⁻¹ galactose). Arabinose utilization is not shown here because arabinose metabolic genes are restored in the subsequent strain described in Figure 4. (d) Consumption of glucose (blue) and xylose (orange) by DLF_GBPC1 in GXA media (no galactose) containing glucose, xylose, and arabinose at a ratio of 11.75:6.25:1 (27.03 g L⁻¹ glucose, 14.38 g L⁻¹ xylose, 2.31 g L⁻¹ arabinose). Arabinose is not consumed in this strain for the same reason noted above.

### Galactose Utilization is Enabled by Altering Glucose Transporter

Having confirmed simultaneous co-utilization of glucose and xylose, we next examined whether strain DLF_GBPC1 could also co-consume galactose and arabinose. We first evaluated galactose utilization in a four-sugar (GXAG) mixture with ratios mimicking lignocellulosic hydrolysate from cornstover (23.5:12.5:2:1).^30^

In four sugar (GXAG) media, the GBP strain DLF_GBPC1 exhibited an unexpected phenotype, preferentially consuming xylose and galactose while severely inhibiting glucose uptake (Figure 2c). Arabinose utilization is not shown in this section because arabinose metabolic genes are restored in the subsequent strain described in Figure 4. We hypothesized that this cross inhibition was mechanistic and arose from competition between glucose and galactose for the GalP permease, which transports galactose as its native substrate as well as glucose. Such competition would limit glucose entry. Consistent with this hypothesis, removing galactose restored efficient glucose–xylose co-utilization (Figure 2d), indicating that GalP mediated transport competition was the primary cause of the observed inhibition.

Based on our hypothesis that galactose competes with glucose for the GalP permease, we replaced the overexpressed *galP* gene cassette in DLF_GBPC1 with alternative glucose transporters, Glf from *Zymomonas mobilis* (DLF_GBPC2) or GlcP from *Bacillus subtilis* (DLF_GBPC3), while retaining the native *galP* copy to maintain galactose uptake (Figure 3a,b).^31,32^ Following strain construction via recombineering,^33^ we screened 10 colonies of each strain for glucose consumption in order to assess potential colony-to-colony variation (Figure S2). From these screenings, single colonies of DLF_GBPC2 and DLF_GBPC3 were selected for further characterization.

**Figure 3.**
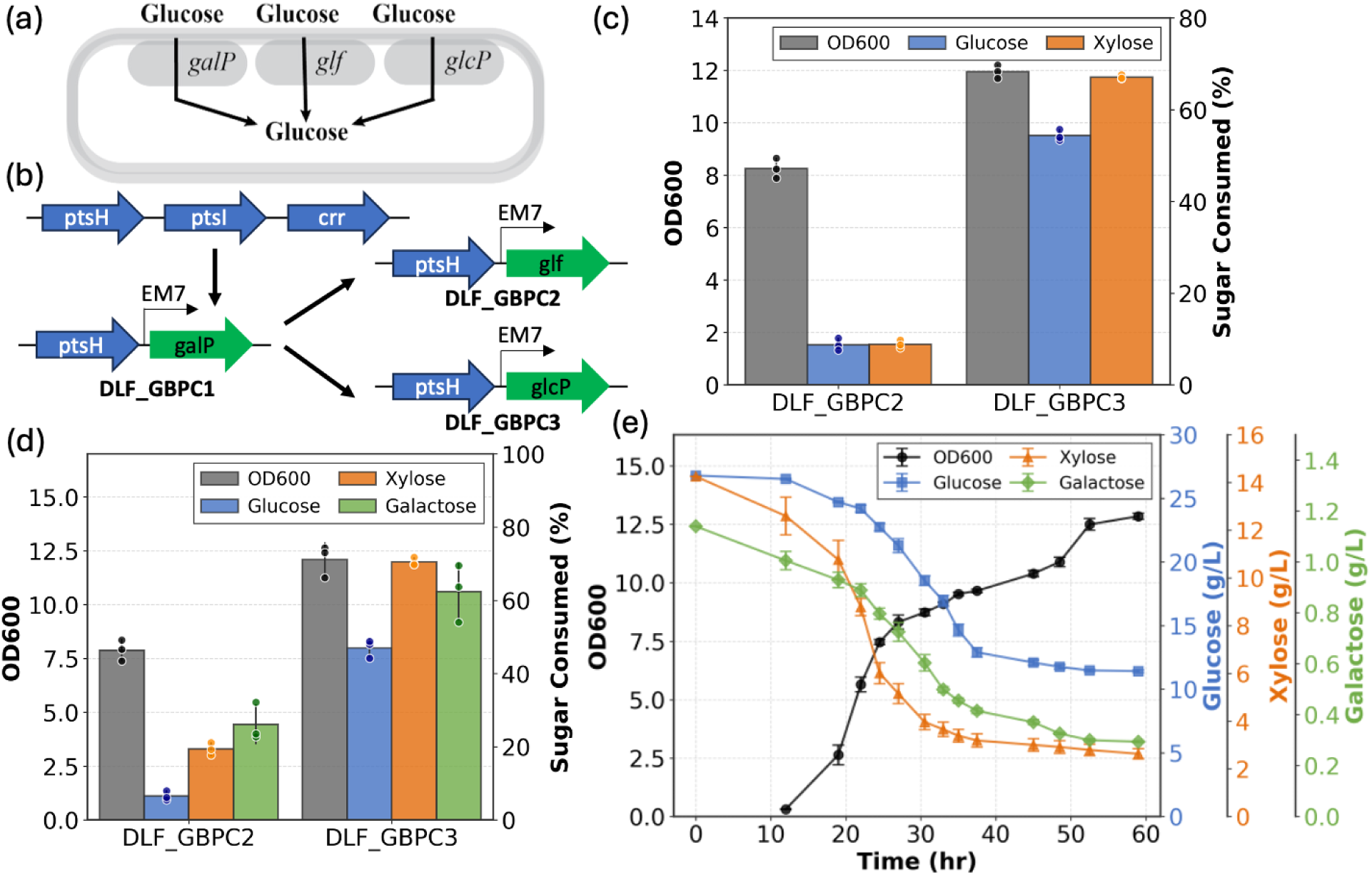
Co-utilization of glucose, xylose, and galactose enabled by altering glucose transport. (a) Schematic of the glucose transport systems evaluated for the GBP architecture. (b) Chromosomal modifications used to construct DLF_GBPC1, and subsequent transporter replacements used to construct DLF_GBPC2 (expressing Glf) and DLF_GBPC3 (expressing GlcP). (c) Co-utilization of glucose (blue) and xylose (orange) by DLF_GBPC2 and DLF_GBPC3 in GXA media containing glucose, xylose, and arabinose at a ratio of 11.75:6.25:1(27.03 g L⁻¹ glucose, 14.38 g L⁻¹ xylose, 2.31 g L⁻¹ arabinose). (d) Co-utilization of glucose (blue), xylose (orange), and galactose (green) by DLF_GBPC2 and DLF_GBPC3 in GXAG media containing glucose, xylose, arabinose, and galactose at a ratio of 23.5:12.5:2:1 (27.03 g L⁻¹ glucose, 14.38 g L⁻¹ xylose, 2.31 g L⁻¹ arabinose, 1.15 g L⁻¹ galactose).(e) Time-course profiles of biomass (black), glucose (blue), xylose (orange), and galactose (green) consumption for DLF_GBPC3 in GXAG media at the same sugar ratio.

**Figure 4.**
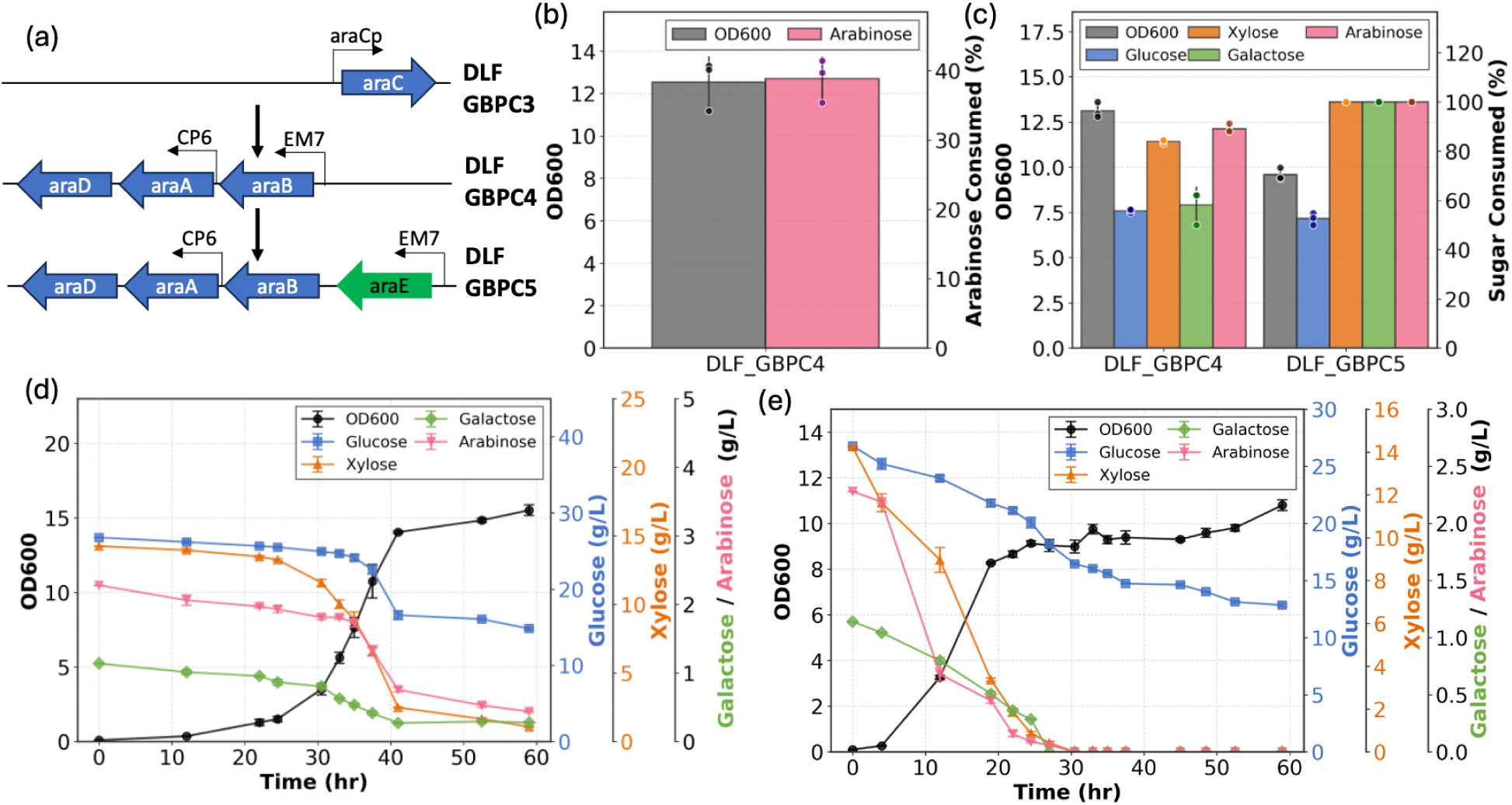
Co-utilization of glucose, xylose, galactose, and arabinose enabled by introducing arabinose metabolic pathway genes. (a) Genetic modifications used to construct DLF_GBPC4 and DLF_GBPC5. DLF_GBPC4 was generated by reintroducing the *araBAD* operon while deleting the *araC* gene. DLF_GBPC5 was derived from DLF_GBPC4 by introducing an additional copy of the *araE* gene. (b) Growth and arabinose consumption by DLF_GBPC4 grown in a minimal media containing arabinose as the sole carbon source. (c) Growth and consumption of glucose, xylose, galactose, and arabinose by DLF_GBPC4 and DLF_GBPC5 in a minimal media containing all four sugars at a ratio of 23.5:12.5:2:1 (27.03 g L⁻¹ glucose, 14.38 g L⁻¹ xylose, 2.31 g L⁻¹ arabinose, 1.15 g L⁻¹ galactose). (d) Time-course profiles of biomass (black), glucose (blue), xylose (orange), arabinose (pink), and galactose (green) consumption for DLF_GBPC4 in GXAG media at the same sugar ratio. Biomass, glucose, and xylose are each plotted on their respective y-axis, while arabinose and galactose are plotted on a shared y-axis. (e) Time-course profiles of biomass (black), glucose (blue), xylose (orange), arabinose (pink), and galactose (green) consumption for DLF_GBPC5 in GXAG media at the same sugar ratio. Biomass, glucose, and xylose are each plotted on their respective y-axis, while arabinose and galactose are plotted on a shared y-axis.

We first evaluated glucose and xylose co-utilization in glucose–xylose–arabinose media lacking galactose using the newly constructed strains (Figure 3c). DLF_GBPC2 (Glf) exhibited reduced consumption of glucose and xylose relative to DLF_GBPC1, whereas DLF_GBPC3 (GlcP) efficiently co-consumed both sugars at levels comparable to the parental strain. We next assessed co-utilization of glucose, xylose, and galactose in glucose–xylose–arabinose–galactose (GXAG) media (Figure 3d). Under these conditions, DLF_GBPC3 simultaneously consumed glucose, xylose, and galactose despite the presence of galactose. These results support the hypothesis that galactose limits glucose uptake in DLF_GBPC1 by competing for GalP, and that replacing GalP-dependent glucose transport with GlcP resolves this bottleneck. Having confirmed restored co-utilization of glucose, xylose, and galactose, we next quantified sugar uptake rates for DLF_GBPC3 in GXAG media (Figure 3e). DLF_GBPC3 consumed glucose at 0.83 g L⁻¹ h⁻¹, xylose at 0.74 g L⁻¹ h⁻¹, and galactose at 0.035 g L⁻¹ h⁻¹. The resulting uptake rate ratios (23.7:21.1:1) closely matched the initial sugar composition (23.5:12.5:1). Although xylose uptake was faster relative to its initial proportion compared with the other sugars, the simultaneous consumption of all three sugars further demonstrates the suitability of this strain for the co-consumption of sugars in lignocellulosic hydrolysates.

### Arabinose Utilization

With this success we turned to the co-consumption of arabinose. The parent *E. coli* strain of the lineage we used to construct the original GBP dependent strains is BW25113, which has a deletion in the arabinose utilization operon *araBAD.*^34^ To restore arabinose metabolism, we reintroduced a functional *araBAD* operon while deleting *araC* in the DLF_GBPC3 background, generating strain DLF_GBPC4 (Figure 4a). We first confirmed that DLF_GBPC4 could efficiently grow on minimal media containing arabinose as the sole carbon source (Figure 4b). Upon confirming restored arabinose catabolism, we then assessed glucose–xylose–galactose–arabinose co-consumption in DLF_GBPC4 (Figure 4c). The strain efficiently utilized all four sugars simultaneously, indicating that deletion of *araC* alleviated reciprocal regulation between the xylose and arabinose pathways.^27,28^ Interestingly, enabling arabinose metabolism also led to higher glucose and xylose consumption in DLF_GBPC4 compared to DLF_GBPC3.

Based on this observation, we explored whether further increasing arabinose uptake could enhance overall sugar co-utilization. To test this, we overexpressed the arabinose transporter *araE* in DLF_GBPC4, generating DLF_GBPC5, following previous reports that araE overexpression improves arabinose assimilation (Figure 4a).^23^ As expected, DLF_GBPC5 showed increased arabinose consumption and completely depleted arabinose from the media. Xylose utilization also increased, consistent with the known promiscuity of the AraE transporter, while glucose consumption remained comparable to DLF_GBPC4. Notably, enhanced arabinose uptake in DLF_GBPC5 also resulted in complete depletion of both xylose and galactose, a metabolic outcome not observed in any prior strain (Figure 4c). Because the regulatory gene araC was deleted in both strains, this effect likely reflects a broader physiological response to increased carbon flux, potentially driven by AraE mediated transport promiscuity or global shifts in cellular energy metabolism. Having confirmed co-utilization of all four sugars, we next quantified sugar uptake rates for DLF_GBPC4 and DLF_GBPC5 in GXAG media (Figure 4d, e). DLF_GBPC4 consumed glucose at 1.0 g L⁻¹ h⁻¹, xylose at 0.96 g L⁻¹ h⁻¹, arabinose at 0.14 g L⁻¹ h⁻¹, and galactose at 0.042 g L⁻¹ h⁻¹. The resulting uptake rate ratios (23.8:22.9:3.33:1) closely matched the initial sugar composition (23.5:12.5:2:1), with the exception that xylose uptake was faster relative to its initial proportion. DLF_GBPC5 consumed glucose at 0.56 g L⁻¹ h⁻¹, xylose at 0.66 g L⁻¹ h⁻¹, arabinose at 0.11 g L⁻¹ h⁻¹, and galactose at 0.042 g L⁻¹ h⁻¹. The corresponding uptake rate ratios (13.3:15.7:2.62:1) again broadly reflected the initial sugar composition (23.5:12.5:2:1), except that glucose uptake was slower than both its initial proportion and the rate observed in DLF_GBPC4. Despite these differences in uptake rates, both strains exhibited simultaneous consumption of all four major sugars present in lignocellulosic hydrolysates.

### Robust Co-utilization

The ability to maintain robust sugar consumption despite variations in feedstock composition is a critical trait for industrial application. To evaluate the robustness of our engineered strains capable of co-consuming all four sugars, DLF_GBPC4 (araBAD restoration) and DLF_GBPC5 (araBAD + araE overexpression), we employed microfermentations to sample a range of lignocellulosic hydrolysate compositions that simulate real world feedstock diversity. We systematically varied the relative concentrations of glucose, xylose, arabinose, and galactose around a center point ratio (23.5:12.5:2:1) to reflect real-world feedstock conditions. The variation range was selected to encompass the typical fluctuations found across different lignocellulosic sources, including agricultural residues and woody biomass.^35–40^

Initially, to evaluate robustness, the concentration of the cellulosic fraction (glucose) was varied from 9 g L⁻¹ to 36 g L⁻¹ while maintaining a constant total sugar concentration of 45 g L⁻¹. These conditions are referred to as G20, G30, G40, G50, G60, G70, and G80, corresponding to glucose concentrations of 9.0, 13.5, 18.0, 22.5, 27.0, 31.5, and 36.0 g L⁻¹, respectively. The remaining sugars, representing the hemicellulosic fraction, were adjusted according to glucose concentration while preserving the same relative ratios as the original GXAG media (12.5:2:1 for xylose:arabinose:galactose). The G60 condition is identical to the original GXAG media and therefore serves as the center point reference for the robustness study. The resulting media compositions are shown in Figure 5a and Table S4.

**Figure 5.**
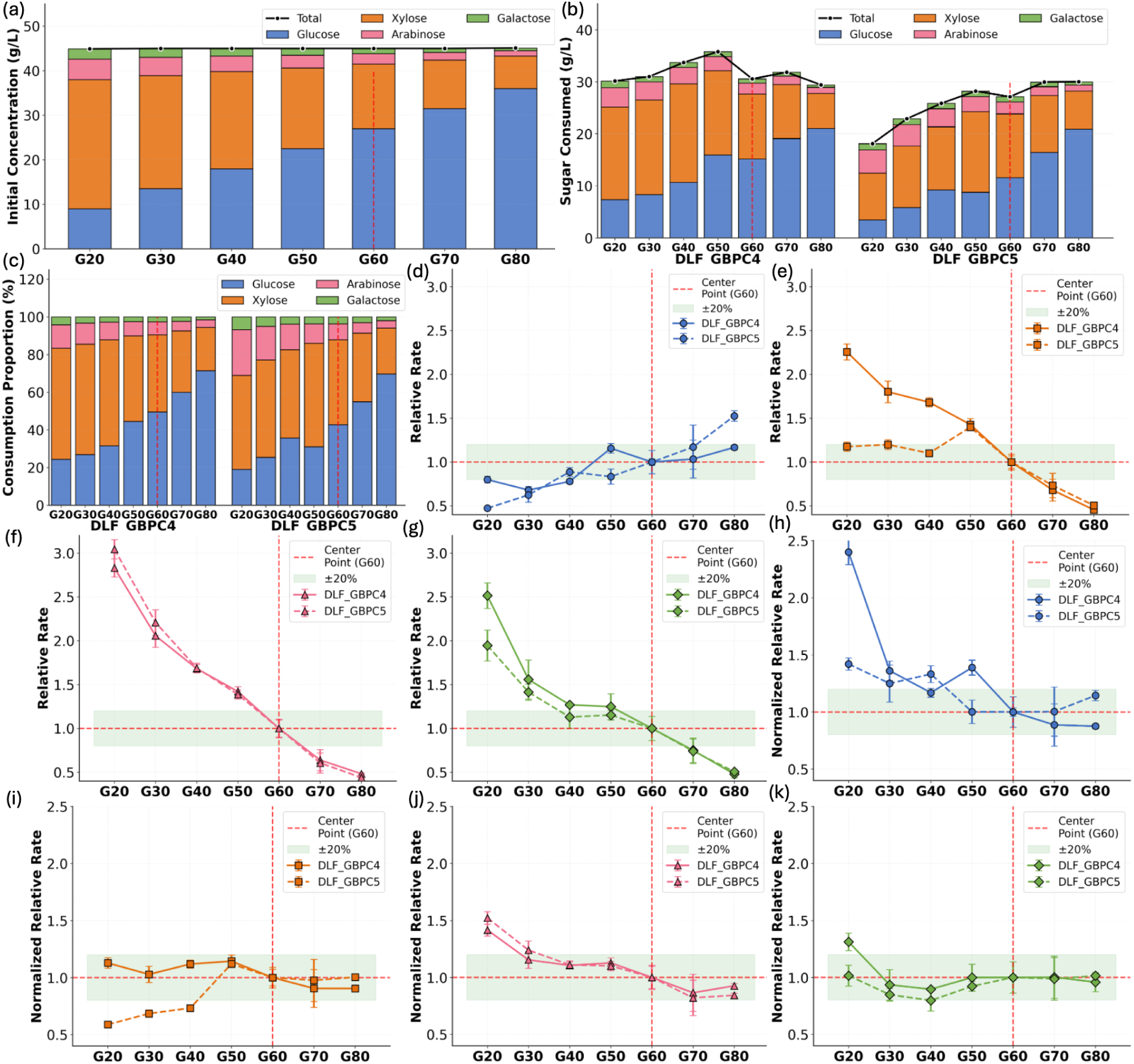
Robustness of engineered GBP strains (DLF_GBPC4 and GBPC5) to variations in feedstock composition. (a) Sugar compositions used to evaluate robustness. The total sugar concentration was held constant (45 g L⁻¹) while initial glucose (G) levels were systematically varied. Concentrations of the remaining sugars (xylose, arabinose, and galactose) were adjusted proportionally to maintain their original ratio in GXAG media (12.5:2:1 relative to glucose), mimicking the compositional heterogeneity of cellulosic fraction in lignocellulosic hydrolysates. The “G60” center point represents a standard ratio of 23.5:12.5:2:1 (27.03 g L⁻¹ glucose, 14.38 g L⁻¹ xylose, 2.31 g L⁻¹ arabinose, 1.15 g L⁻¹ galactose). (b) Total sugar consumed (g L⁻¹) after 50 hrs by DLF_GBPC4 (araBAD restoration) and DLF_GBPC5 (araBAD + araE overexpression). (c) Percentage of each sugar in total consumption across conditions by DLF_GBPC4 and DLF_GBPC5. (d–g) Relative sugar uptake rates for strains DLF_GBPC4 (solid lines) and DLF_GBPC5 (dashed lines) across varying glucose concentrations (G20–G80) for glucose (d), xylose (e), arabinose (f), and galactose (g). Rates are shown relative to G60 center point (red line). The green shaded region indicates the ±20% threshold. (h–k) Normalized relative sugar uptake rates for strains DLF_GBPC4 (solid lines) and DLF_GBPC5 (dashed lines) across varying glucose concentrations (G20–G80) for glucose (h), xylose (i), arabinose (j), and galactose (k). Relative rates from panels (d–g) are further normalized by their respective initial sugar concentrations to account for differences in initial sugar levels (rate per initial concentration).

The amount of each sugar consumed, as well as total sugar consumption, by DLF_GBPC4 and DLF_GBPC5 is shown in Figure 5b. DLF_GBPC4 consumed approximately 30 g L⁻¹ total sugars across all conditions. In contrast, DLF_GBPC5 exhibited conditiondependent variation, consuming approximately 18 g L⁻¹ in G20 media and approximately 30 g L⁻¹ in G80 media. In both strains, glucose consumption increased with increasing initial glucose concentration. To better visualize changes in sugar utilization patterns, sugar consumption was also expressed as the fraction of each sugar relative to total sugar consumed (Figure 5c). Interestingly, as the availability of individual sugar components varied across the G20–G80 series, both strains adjusted their consumption profiles accordingly.

We next examined sugar uptake rates for both strains relative to the G60 condition, which corresponds to the original GXAG media used in prior shake flask experiments (Figure 5d–g). Glucose uptake rates were similar across conditions, whereas uptake rates for xylose, arabinose, and galactose decreased as their respective initial concentrations decreased. To account for differences in starting concentrations, uptake rates were next divided by the initial concentration of each sugar (Figure 5h–k). With uptake rates expressed relative to initial sugar concentrations, uptake rates were comparable across all conditions, further indicating that the strains adjust their sugar consumption rates in response to feedstock composition.

We next performed a similar set of studies varying the composition of the hemicellulosic fraction of the sugar mixture using DLF_GBPC4, which previously showed greater robustness across conditions compared to DLF_GBPC5. In this case, glucose concentration was held constant while the ratios of hemicellulosic sugars were varied to simulate highly heterogeneous hemicellulose profiles (Figure 6a). To achieve this, low- and high-glucose background conditions, G20 and G70 media, were selected, and their hemicellulosic compositions were systematically varied to generate the H1–H14 media series, as detailed in Table S4.

**Figure 6.**
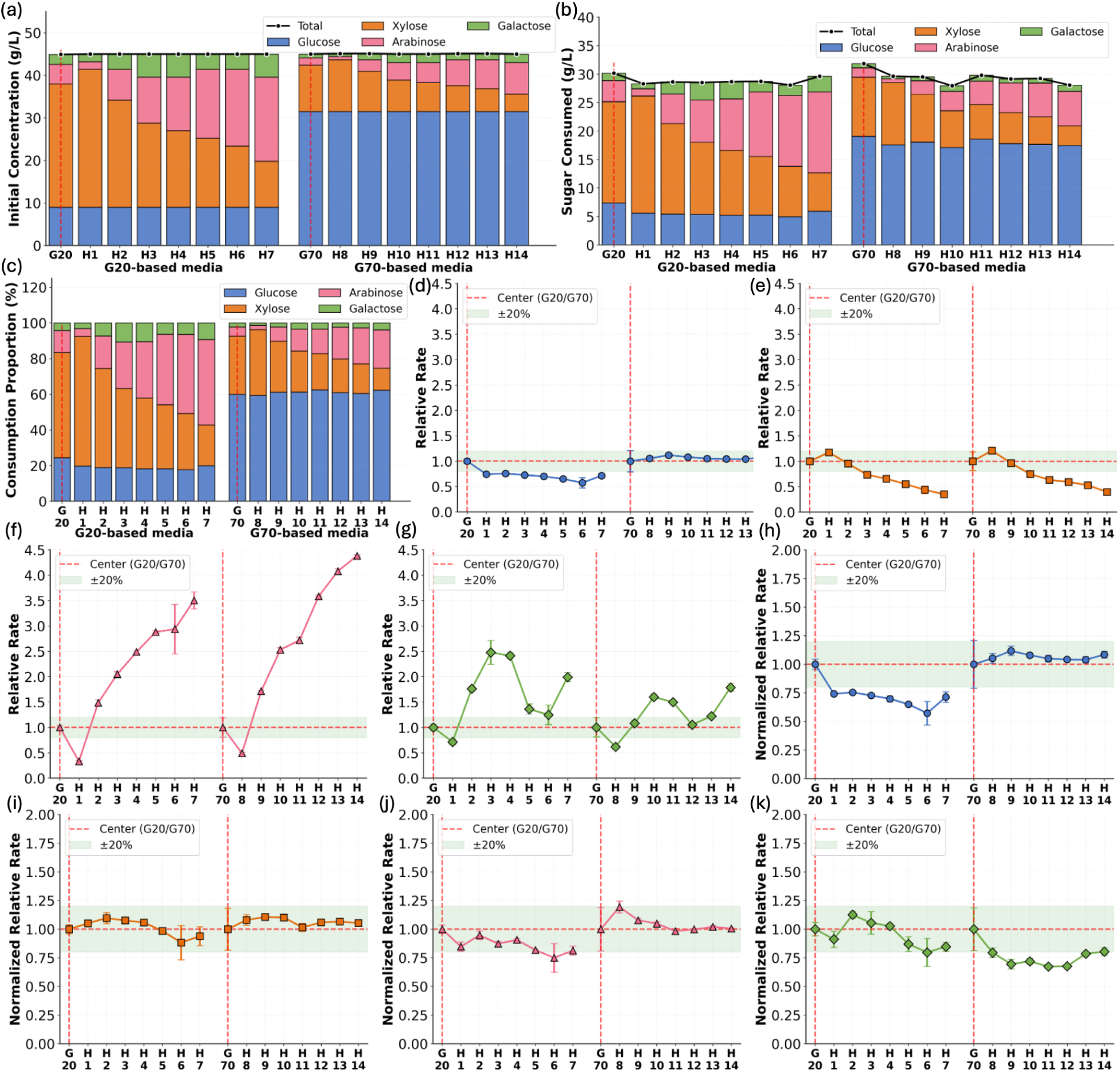
Robustness of DLF_GBPC4 to variations in hemicellulose sugar composition. (a) Initial sugar concentrations across varied hemicellulose compositions. Sugar composition variations simulating highly variable hemicellulose profiles while maintaining a fixed glucose baseline. The G20-based and G70-based media series (Figure 5) represent low- and high-glucose backgrounds, respectively. Within each background, defined ratios of xylose, arabinose, and galactose were applied (H1–H14). (b) Total sugar consumed (g L⁻¹) after 50 hrs by DLF_GBPC4 demonstrating consistent simultaneous uptake of sugars despite variation in hemicellulose composition (c) Percentage of each sugar in total consumption across conditions by DLF_GBPC4. (d–g) Relative consumption rates across varying hemicellulose composition for DLF_GBPC4 for glucose (d), xylose (e), arabinose (f), and galactose (g). Rates are shown relative to the corresponding G20 or G70 reference media (red lines) used to generate each hemicellulose composition. The green shaded region indicates the ±20% threshold. (h–k) Normalized relative sugar uptake rates for strains DLF_GBPC4 for glucose (h), xylose (i), arabinose (j), and galactose (k). Relative rates from panels (d–g) are further normalized by their respective initial sugar concentrations to account for differences in initial sugar levels (rate per initial concentration).

Across all conditions, DLF_GBPC4 consistently consumed ∼30 g L⁻¹ total sugars while modulating the consumption of individual sugars according to their availability, demonstrating robustness across substantial variation in hemicellulosic composition and C5:C6 sugar balance (Figure 6b,c). Consistent with the glucose concentration being held constant, glucose uptake rates were similar across conditions, whereas uptake rates for xylose, arabinose, and galactose decreased as their respective initial concentrations decreased (Figure 6d–g). When uptake rates were expressed relative to initial sugar concentrations, rates were consistent across all conditions, confirming that DLF_GBPC4 with gluconate-bypass metabolism, which is decoupled from native regulatory feedback, maintains robust performance across diverse feedstock compositions (Figure 6h–k).

## Discussion

Our study demonstrates that strategically rewiring central carbon metabolism through a Gluconate Bypass (GBP) enables the simultaneous co-consumption of all four major lignocellulosic sugars (glucose, xylose, arabinose, and galactose). More importantly, strains with the GBP metabolism adapt to varying lignocellulosic sugar compositions by adjusting individual sugar uptake rates according to sugar availability, demonstrating robustness. Here, robustness is defined as the ability to maintain stable performance across varying feedstock compositions. The central innovation lies in decoupling glucose consumption from the native glycolytic network and its regulatory feedback, rather than operating within the constraints imposed by these regulations as in previous approaches. By fully decoupling sugar metabolism from native regulation, strains with GBP metabolism effectively circumvent carbon catabolite repression imposed by glucose uptake and glycolytic regulation. This decoupling not only enables co-consumption of all four sugars but also allows robust performance across a wide range of lignocellulosic sugar compositions, which is critical for industrial scale-up of bioprocesses using lignocellulosic feedstocks.

By converting glucose to gluconate, which subsequently enters the pentose phosphate pathway (PPP) and the Entner–Doudoroff (ED) pathway, the GBP architecture channels all major sugars into shared metabolic routes. While this architecture initially enabled co-consumption of glucose and xylose, additional optimizations were needed to address sugar-specific challenges and achieve simultaneous utilization of all four major lignocellulosic sugars. First, we identified an unexpected inhibition of glucose uptake caused by competition with galactose for the GalP permease. This limitation was resolved by replacing GalP with the glucose specific transporter GlcP, which restored co-utilization of glucose, xylose, and galactose. Second, we enabled simultaneous consumption of all four major lignocellulosic sugars by reintroducing the *araBAD* operon, which was absent in the initial strain with GBP metabolism.

The result of this systematic engineering is the first reported instance of an *E. coli* strain achieving robust co-utilization of glucose, xylose, arabinose, and galactose. Across all conditions tested, strain DLF_GBPC4 simultaneously consumed all four sugars while regulating individual sugar uptake rates in response to the availability of each component. The robustness of these strains across a sampled range of lignocellulosic hydrolysate compositions that simulate real world feedstock diversity supports their use as industrially-ready chassis. Unlike previous approaches that rely on complex co-culture strategies, this single-strain solution simplifies the bioprocess by eliminating the need for external process control and allowing the cell’s own metabolism to regulate sugar utilization, providing a scalable platform for sustainable lignocellulose-based fermentation. Moving forward, this robust GBP metabolism provides a powerful foundation for solving the next generation of biorefining challenges. The platform can be further modified to incorporate other minor lignocellulosic components such as mannose, acetate and cellobiose.^41^ The decoupled architecture is ideally suited for integration with advanced bioprocessing strategies, such as two-stage dynamic metabolic control,^42–44^ although issues with common inhibitory compounds like lignin-derived toxins (furfurols) in real-world hydrolysates must be addressed in future work.^45,46^

In conclusion, our approach of engineering the central carbon metabolism to bypass glycolysis fundamentally shifts the paradigm from mitigating repression to decoupling metabolic regulation. This technology offers a robust and scalable platform critical for realizing the full potential of lignocellulosic biomass in sustainable biomanufacturing.

## Author contributions

U. Yano constructed strains, performed microfermentations and growth curve studies as well as analytical analyses. P. Sarkar performed microfermentations. M.D. Lynch designed experiments. All authors analyzed results, wrote, revised and edited the manuscript.

## Acknowledgements

We would like to acknowledge the following support: DOE EERE grant #EE000756, NSF # 2350533, U. Yano was supported by the Takenaka Scholarship Foundation.

## Materials & Methods

### Reagents and media

Unless otherwise specified, all materials and reagents were purchased from Sigma-Aldrich (St. Louis, MO, USA). Luria Broth (Lennox formulation) was used for routine strain and plasmid propagation and construction. Working antibiotic concentrations were as follows: kanamycin (35 μg/mL), chloramphenicol (35 μg/mL), zeocin (50 μg/mL), gentamicin (25 μg/mL), blasticidin (100 μg/mL), and spectinomycin (50 μg/mL). SM10++, SM10^1/2,½^ , SM10 media were prepared according to previously reported protocols^1,2^ with modification of adding sugars. Glucose-xylose (GX) media was formulated to have the final concentration of glucose to be 30 g L⁻¹ and xylose to be 15 g L⁻¹. Glucose-xylose-arabinose (GXA) media was made to have the final concentration of glucose to be 27.03 g L⁻¹, xylose to be 14.38 g L⁻¹, and arabinose to be 2.31 g L⁻¹. Glucose-xylose-arabinose (GXAG) media was made to have the final concentration of glucose to be 27.03 g L⁻¹, xylose to be 14.38 g L⁻¹, arabinose to be 2.31 g L⁻¹, and galactose to be 1.15 g L⁻¹. SM10 arabinose media was formulated to have the final concentration of 45 g L⁻¹ just as regular SM10 media would be made with glucose.

### Strains

All host strains were constructed using standard recombineering methods as previously described.^3^ A complete list of strains is provided in Supplementary Table S1. The recombineering plasmid pSIM5 was kindly provided by Donald Court (NCI; https://redrecombineering.ncifcrf.gov/court-lab.html).^4^ Recombinant strains were verified by PCR and DNA sequencing (Azenta Life Sciences, MA).

### Shake flask experiment

A 5 mL Luria Broth (LB) culture was inoculated from glycerol stock and incubated overnight at 37 °C with shaking. The following day, 1% (v/v) of the overnight culture was transferred into a 250 mL baffled shake flask containing 25 mL SM10++ media with appropriate sugar mixtures and incubated at 37 °C until the culture reached an OD_600_ of 6–10. Subsequently, 1% (v/v) of this culture was used to inoculate 25 mL of SM10 media (containing half the concentrations of casamino acids and yeast extract present in SM10++), and incubation was continued under identical conditions until OD_600_ reached 6–10. Finally, 1% (v/v) of this culture was used to inoculate SM10 media lacking casamino acids and yeast extract. For the end-point study, samples were taken at 50 hrs. For the time course samples were taken multiple times during the growth. Sugar uptake rates were calculated by determining the slope of the exponential phase using a sliding window of four consecutive data points and selecting the maximal rate across all possible windows.

### Robustness study

The same adaptation protocol was followed and DLF_GBPC4 and DLF_GBPC5 were adapted to original GXAG media with all four sugars at a ratio of 23.5:12.5:2:1. This adapted cell was used to inoculate SM10 media lacking casamino acids and yeast extract with varied concentration of glucose, xylose, arabinose, and galactose in a 96-well plate. These plates were incubated at 37°C with plate covers, shaking at 300 rpm at a 50 mm orbit following previous microfermentation protocol for 50 hrs.^1,5^

### Analytical methods

#### Glucose, xylose, arabinose, and galactose quantification

All sugar concentrations in shake flask and microfermentation samples were measured using a UPLC system (Acquity H-Class, Waters Corp., MA, USA) equipped with a 2414 Refractive Index (RI) detector. For glucose and arabinose, chromatographic separation was achieved on a Rezex ROA-Organic Acid H⁺ (8%) column (300 × 7.8 mm; Cat. no. 00H-0138-K0, Phenomenex, Inc., CA, USA) maintained at 50 °C. The mobile phase consisted of 5 mM sulfuric acid, delivered isocratically at 0.5 mL min⁻¹ for 35 min. The injection volume was 10 μL. To maintain linearity within the analytical range, samples were diluted 5-fold. For xylose and galactose, chromatographic separation was achieved on a Restek Ultra Amino column (150 x 4.6 mm, 3 μm; Cat. no. 9107365, Restek Corporation, Bellefonte, PA) maintained at 35 °C. The mobile phase consisted of Water:acetonitrile (25:75), delivered isocratically at 0.8 mL min⁻¹ for 20 min. The injection volume was 10 μL.

