## Supplemental Materials for "Decoupling Central Metabolism from Catabolite Repression Enables Robust Cellulosic Sugar Co-consumption in *E. coli*"

### Supplementary Materials

**Table S1: Plasmids & Strains used in this study**

| Plasmid | Purpose | Origin | Marker | Addgene | Source |
|---|---|---|---|---|---|
| pSIM5 | Genome Editing | pSC101ts | Cm | NA | Court Lab |
| pSMART-GFPuv | GFPuv expression | colE1 | Kan | 65822 | |

| Strain | Purpose | Genotype | Source |
|---|---|---|---|
| DLF_Z0025 | Dynamic Control | F-, λ-, Δ(araD-araB)567, lacZ4787(del)(::rrnB-3) , rph-1, Δ(rhaD-rhaB)568, hsdR514, ΔackA-pta, ΔpoxB, ΔpflB, ΔldhA, ΔadhE, ΔiclR, ΔarcA, ΔsspB::frt, Δcas3::tm-ugpb-sspB-pro-casA | [1] |
| DLF_S0025 | Dynamic Control | F-, λ-, Δ(araD-araB)567, lacZ4787(del)(::rrnB-3) , rph-1, Δ(rhaD-rhaB)568, hsdR514, ΔackA-pta, ΔpoxB, ΔpflB, ΔldhA, ΔadhE, ΔiclR, ΔarcA, ΔsspB::frt, Δcas3::tm-ugpb-sspB-pro-casA | [2] |
| DLF_0286 | Dynamic Control | DLF_Z0025, ΔptsG::proCp-glk, galPp::proCp | (Yano et al. 2025) |
| DLF_GBPC1 | gluconate bypass | DLF_S0025, ΔptsI/crr::EM7-galP, ΔgntR::EM7-gdh-gnl | (Yano et al. 2025) |
| DLF_GBPC2 | glf | DLF_S0025, ΔptsI/crr::EM7-glf, ΔgntR::EM7-gdh-gnl | this study |
| DLF_GBPC3 | glcP | DLF_S0025, ΔptsI/crr::EM7-glcP, ΔgntR::EM7-gdh-gnl | this study |
| DLF_GBPC4 | glcP, araBAD | DLF_GBPC3, ΔaraC::EM7-araB-CP6-araA-araD | this study |
| DLF_GBPC5 | glcP,araE, araBAD | DLF_GBPC3, ΔaraC::EM7-araE-araB-CP6-araA-araD | this study |

**Table S2:** Synthetic DNA utilized for strain construction.

| **ΔptsI/crr-glf-specR** |
| --- |
| TGTGACTATCTCCGCAGAAGGCGAAGACGAGCAGAAAGCGGTTGAACATCTGGTTAAACTGATGGCGGAACTC<br>GAGtaaTTTCCCGGGACCGTGTTGACAATTAATCATCGGCATAGTATATCGGCATAGTATAATACGACAACAACATT<br>ATACAACGACCCCAGAGGGACAAGATTAAGGAGGTAATTTATGAGCAGCGAAAGCTCACAAGGGCTGGTTACA<br>CGCTTGGCCCTGATTGCGGCAATTGGAGGACTTTTATTCGGTTACGATAGCGCAGTCATTGCGGCTATTGGCAC<br>ACCCGTCGACATCCATTTCATCGCTCCGCGCCACCTTAGCGCTACGGCGGCTGCATCGTTGAGTGGCATGGTT<br>GTGGTAGCCGTTCTGGTAGGTTGCGTGACCGGTTCACTGTTGTCAGGCTGGATCGGGATTCGTTTTGGGCGT<br>CGCGGTGGCTTACTGATGTCATCTATCTGCTTTGTTGCCGCCGGGTTTGGGGCAGCTCTTACGGAGAAGCTGT<br>TTGGTACTGGCGGTTCTGCATTGCAGATCTTTTGTTTCTTCCGCTTCCTTGCAGGGCTTGGCATCGGAGTAGTC<br>TCAACGCTGACTCCCACGTATATTGCTGAAATCGCGCCTCCTGACAAACGTGGTCAAATGGTGTCTGGCCAAC<br>AGATGGCAATCGTAACCGGTGCATTGACGGGGTATATTTTCACCTGGCTTTTAGCACACTTTGGTAGTATTGATT<br>GGGTTAACGCGTCTGGATGGTGCTGGTCACCCGCATCAGAAGGGCTTATTGGTATTGCCTTTCTTCTTTTATTA<br>CTGACTGCGCCTGATACACCACATTGGTTAGTCATGAAGGGGCGTCATTCGGAGGCAAGCAAGATTCTGGCAC<br>GTCTTGAACCTCAGGCTGACCCAAATTTGACAATTCAAAAGATTAAGGCCGGTTTCGATAAAGCTATGGATAAGA<br>GCAGCGCGGGTTTATTCGCATTCGGTATTACTGTAGTGTTTGCAGGAGTGTCCGTAGCGGCATTCCAGCAGCTT<br>GTCGGCATCAACGCCGTGCTGTACTACGCTCCCCAGATGTTCCAAAACTTAGGCTTCGGAGCTGATACCGCGT<br>TACTGCAAACAATTTCCATTGGGGTCGTTAATTTCATTTTCACAATGATCGCGTCGCGCGTCGTGGACCGCTTT<br>GGCCGCAAGCCATTGCTGATTTGGGGTGCACTGGGTATGGCCGCTATGATGGCTGTACTTGGGTGCTGTTTTT<br>GGTTTAAGGTGGGGGGTGTGTTACCTTTGGCCTCCGTATTACTTTATATTGCAGTATTCGGGATGTCATGGGGT<br>CCCGTCTGCTGGGTCGTTCTTAGCGAAATGTTCCCTAGCTCGATCAAAGGAGCTGCAATGCCCATCGCAGTAA<br>CTGGTCAGTGGCTGGCCAACATCCTTGTAAACTTCCTTTTCAAGGTTGCCGACGGTAGCCCTGCTTTAAATCAA<br>ACGTTTAACCACGGGTTTTCCTACTTGGTGTTTGCGGCACTGAGCATCCTTGGAGGTCTGATTGTCGCGCGTT<br>TTGTCCCGGAAACTAAAGGGCGCTCTTTAGACGAAATTGAAGAAATGTGGCGTAGCCAGAAGTGAAGATCGGC<br>ACGTAAGAGGTTCCAACTTTCACCATAATGAAATAAGATCACTACCGGGCGTATTTTTTGAGTTATCGAGATTTTC<br>AGGAGCTAAGGAAGCTAAAATGAGGGAAGCGGTGATCGCCGAAGTATCGACTCAACTATCAGAGGTAGTTGGC<br>GTCATCGAGCGCCATCTCGAACCGACGTTGCTGGCCGTACATTTGTACGGCTCCGCAGTGGATGGCGGCCTG<br>AAGCCACACAGTGATATTGATTTGCTGGTTACGGTGACCGTAAGGCTTGATGAAACAACGCGGCGAGCTTTGAT<br>CAACGACCTTTTGGAAACTTCGGCTTCCCCTGGAGAGAGCGAGATTCTCCGCGCTGTAGAAGTCACCATTGTT<br>GTGCACGACGACATCATTCCGTGGCGTTATCCAGCTAAGCGCGAACTGCAATTTGGAGAATGGCAGCGCAATG<br>ACATTCTTGCAGGTATCTTCGAGCCAGCCACGATCGACATTGATCTGGCTATCTTGCTGACAAAAGCAAGAGAA<br>CATAGCGTTGCCTTGGTAGGTCCAGCGGCGGAGGAACTCTTTGATCCGGTTCCTGAACAGGATCTATTTGAGG<br>CGCTAAATGAAACCTTAACGCTATGGAACTCGCCGCCCGACTGGGCTGGCGATGAGCGAAATGTAGTGCTTAC<br>GTTGTCCCGCATTTGGTACAGCGCAGTAACCGGCAAAATCGCGCCGAAGGATGTCGCTGCCGACTGGGCAAT<br>GGAGCGCCTGCCGGCCCAGTATCAGCCCGTCATACTTGAAGCTAGACAGGCTTATCTTGGACAAGAAGAAGAT<br>CGCTTGGCCTCGCGCGCAGATCAGTTGGAAGAATTTGTCCACTACGTGAAAGGCGAGATCACCAAGGTAGTC<br>GGCAAATAATTTTGCCGCAGTTTATGCTTCCGCCAGCGGCGGCAAAATCAATTCATCGCTCTCATGCTGCTGGG<br>TGTAGCGCATCACTTCCAGTACGCGCAACCCCGCTCGGTGCACTGCATCGGTTAACGCCTTCCCTTTCAGCAA<br>GCCACTGATGAGCTGAGCACAAAA |
| **ΔptsI/crr-glcP-ampR** |
| TAAAAGAAGCTAAGGGCTTCACTTCTGAAATTACTGTGACTTCCAACGGCAAAAGCGCCAGCGCGAAAAGCCT<br>GTTTAAACTGCAGACTCTGGGCCTGACTCAAGGTACCGTTGTGACTATCTCCGCAGAAGGCGAAGACGAGCA<br>GAAAGCGGTTGAACATCTGGTTAAACTGATGGCGGAACTCGAGtaaTTTCCCGGGACCGTGTTGACAATTAATC<br>ATCGGCATAGTATATCGGCATAGTATAATACGACAAGGTGAGGAACTAAACCATGTTGCGTGGAACCTATCTTTTT<br>GGGTATGCCTTCTTTTTCACGGTAGGCATTATCCATATTTCTACAGGGTCATTGACCCCGTTTCTTCTGGAAGCT |

TTCAATAAAACAACAGATGATATTAGTGTAATCATCTTCTTTCAATTTACAGGTTTTCTTAGCGGTGTGCTTATTGC
GCCACTTATGATCAAAAAGTATAGTCACTTTCGCACATTAACACTTGCTCTTACAATTATGTTGGTTGCACTTTCA
ATTTTCTTTCTTACGAAAGATTGGTACTACATTATCGTAATGGCATTTCTGTTGGGTTACGGCGCGGGCACCCTG
GAGACCACGGTAGGTTCGTTTGTAATTGCTAATTTTGAGAGCAATGCGGAAAAGATGTCCAAACTTGAAGTCCT
TTTCGGACTGGGTGCCCTGTCTTTCCCACTTCTGATCAACTCGTTTATCGACATCAATAATTGGTTTTTGCCCTA
CTACTGCATCTTTACCTTCCTTTTCGTGCTTTTCGTGGGATGGTTGATTTTTCTTTCGAAGAACCGCGAATATGC
GAAGAACGCCAATCAGCAGGTGACGTTCCCAGACGGGGGGGCATTTCAGTACTTCATTGGAGACCGCAAAAA
ATCTAAACAATTAGGTTTTTTCGTGTTCTTTGCCTTCTTGTATGCGGGGATCGAGACAAACTTCGCGAATTTTCT
GCCTTCGATTATGATCAACCAGGATAACGAGCAAATCTCTCTGATTTCTGTCTCGTTCTTTTGGGTCGGAATCAT
TATCGGCCGCATCCTGATCGGATTTGTGTCCCGCCGTCTGGATTTCAGCAAATATTTATTATTTAGCTGCAGTTG
CTTAATCGTGCTGTTAATCGCATTTAGTTACATTAGCAACCCTATTCTTCAGTTATCGGGCACATTCCTTATCGGC
CTGAGTATTGCCGGCATCTTCCCCATTGCTCTTACTCTGGCTTCGATTATCATTCAGAAATACGTGGACGAAGTC
ACCAGTCTGTTTATCGCTTCCGCGTCCTTTGGCGGTGCAATTATCTCTTTTCTTATCGGTTGGAGCTTGAACCAA
GATACCATTTTGCTGACAATGGGCATCTTTACCACTATGGCTGTGATCTTGGTTGGTATTAGCGTAAAAATCCGT
CGTACGAAAACAGAAGACCCCATTTCATTAGAGAATAAGGCATCAAAAACCCAATGAAGATCGGCACGTAAGAG
GTTCCAACTTTCACCATAATGAAATAAGATCACTACCGGGCGTATTTTTTGAGTTATCGAGATTTTCAGGAGCTAA
GGAAGCTAAAATGAGTATTCAACATTTCCGTGTCGCCCTTATTCCCTTTTTTGCGGCATTTTGCCTTCCTGTTTTT
GCTCACCCAGAAACGCTGGTGAAAGTAAAAGATGCTGAAGATCAGTTGGGTGCACGAGTGGGTTACATCGAAC
TGGATCTCAACAGCGGTAAGATCCTTGAGAGTTTACGCCCCGAAGAACGTTTTCCAATGATGAGCACTTTTAAA
GTTCTGCTATGTGGCGCGGTATTATCCCGTATTGACGCCGGGCAAGAGCAACTCGGTCGCCGCATACACTATTC
TCAGAATGACTTGGTTGAGTACTCACCAGTCACAGAAAAGCATCTCACGGATGGCATGACAGTAAGAGAATTAT
GCAGTGCTGCCATAACCATGAGTGATAACACTGCGGCCAACTTACTTCTGGCAACGATCGGAGGACCGAAGGA
GCTAACCGCTTTTTTGCACAACATGGGGGATCATGTAACTCGCCTTGATCGTTGGGAACCGGAGCTGAATGAA
GCCATACCAAACGACGAGCGTGACACCACGATGCCTGTAGCAATGGCAACAACGTTGCGCAAACTATTAACTG
GCGAACTACTTACTCTAGCTTCCCGGCAACAATTAATAGACTGGATGGAGGCGGATAAAGTTGCAGGATCACTT
CTGCGCTCGGCCCTCCCGGCTGGCTGGTTTATTGCTGATAAATCTGGAGCCGGTGAGCGTGGGTCTCGCGGT
ATCATTGCAGCACTGGGGCCAGATGGTAAGCCCTCCCGCATCGTAGTTATCTACACGACGGGGAGTCAGGCAA
CTATGGATGAACGAAATAGACAGATCGCTGAGATAGGTGCCTCACTGATTAAGCATTGGTAATTTTGCCGCAGTT
TATGCTTCCGCCAGCGGCGGCAAAATCAATTCATCGCTCTCATGCTGCTGGGTGTAGCGCATCACTTCCAGTAC
GCGCAACCCCGCTCGGTGCACTGCATCGGTTAACGCCTTCCCTTTCAGCAAGCCACTGATGAGCTGAGCACA
AAACAGGTCGCCAGTCCCTTTCAGGTCGGTTTTTACCCGTGAATGGGAAATGACATTC

**ΔaraC-EM7-bsdR-araA**

GGCGTGTCGGAGGAGGGCTTTTTCGTACCTTCAACCACTTCACCGGTTTCGATGCTAACCACGACCATATCGT
CAGCGGTCATGACGCTGTAATCGACGCCGGAAGGTTTGATCACAAAGACGCCGCGCTCGCGATCAACGGCGC
TGACGTTGCCCCATGTGAGCGTGACCAGGTTGTGTTTTGGCAGCGCCAGGTTGGCTTCTAATACCTGGCGTTT
GAGATCTTCTAACATGTTGACTCCTTCGTGCCGGATTAGCGACGAAACCCGTAATACACTTCGTTCCAGCGCAG
CGCGTCTTTAAACGCTGGCAGGCGTGTGTCGTTATCAATCACCGTGATTTCAATGTCGTGCATCTCGGCGAATT
GGCGCATATCGTTGAGGTTCAGTGCATGGCTGAAGACGGTATGGTGCGCGCCACCAGCGAGGATCCACGCTT
CGGAAGCAGTTGGCAGATCCGGTTGCGCTTTCCACAGCGCATTCGCCACCGGCAGTTTCGGCAGGGAGTGC
GGTGTTTTCACCGTGTCGATGCAGTTAACCAGTAGACGGTAACGATCGCCGAGATCAATCAAGCTGGCGACAA
TCGCTGGGCCGGTTTGGGTATTGAAGATCAGGCGGGCAGGATCGTCCTTACCACCAATACCGAGATGCTGAAC
GTCGAGGATCGGTTTCTCTTCTGCGGCGATCGACGGGCAGACTTCCAGCATATGGGAGCCGAGCACCAGGTC
ATTACCTTTCTCGAAGTGATAGGTGTAGTCCTCCATAAAGGAAGTACCACCCTGCAGACCGGTTGACATCACCT
TCATGATGCGAAGCAGGGCGGCAGTTTTCCAGTCGCCTTCGCCCGCAAAGCCGTAACCCTGCTGCATCAGAC
GCTGTACGGCCAGACCAGGAAGCTGTTTCAGACCGTGCAAATCTTCAAAGGTGGTGGTGAACGCGTGGAAGC
CACCTTGTTCCAGGAAACGCTTCATCCCCAGCTCAATACGCGCCGCTTCCAGCACGTTCTGTCGTTTTTTGCC
GTGGATTTGTGTGGCAGGCGTCATGGTGTAGCAGCTTTCGTACTCATCGACCAGCGCGTTAACATCGCCGTCG
CTGATGGAGTTCACCACCTGCACCAGATCGCCAACCGCCCAGGTATTGACGGAGAAACCGAACTTGATCTGTG
CGGCAACTTTATCGCCATCGGTGACCGCCACTTCACGCATGTTATCGCCAAATCGGCAGACTTTCAGATGACG
GGTATCCTGTTTAGAGACCGCCTGACGCATCCAGGAGCCGATACGCTCATGGGCTTGTTTATCCTGCCAGTGA
CCGGTAACCACGGCATGTTGCTGACGCATACGCGCGCCAATGAAGCCGAACTCGCGACCGCCATGTGCAGTC

TGGTTCAGGTTCATAAAGTCCATATCGATACTGTCCCACGGCAGCGCCGCGTTGAACTGGGTGTGGAATTGCA
GCAACGGTTTGTTGAGCATGGTCAGGCCGTTGATCCACATTTTGGCCGGGGAGAAGGTGTGCAGCCACACCA
CCAGACCAGCGCAACGATCGTCGTAATTCGCGTCGCGGCAAATAGCGGTGATTTCATCCGGCGTGGTGCCCA
GCGGTTTCAACACCAGTTTGCAGGGCAGTTTCGCTTCCGTATTCAGCGCATTAACGACGTGCTCGGCATGTTG
GGTGACCTGACGCAGGGTTTCCGGGCCATACAGATGCTGGCTGCCAATGACAAACCACACTTCATAATTATCAA
AAATCGTCATTATCGTGTCCTTACTTTAGTTCCTGGTGTACTTGAGGGGGATGAGTTCCTCAATGGTGGTTTTGA
CCAGCTTGCCATTCATCTCAATGAGCACAAAGCAGTCAGGAGCATAGTCAGAGATGAGCTCTCTGCACATGCC
ACAGGGGCTGACCACCCTGATGGATCTGTCCACCTCATCAGAGTAGGGGTGCCTGACAGCCACAATGGTGTC
AAAGTCCTTCTGCCCGTTGCTCACAGCAGACCCAATGGCAATGGCTTCAGCACAGACAGTGACCCTGCCAATG
TAGGCCTCAATGTGGACAGCAGAGATGATCTCCCCAGTCTTGGTCCTGATGGCCGCCCCGACATGGTGCTTGT
TGTCCTCATAGAGCATGGTGATCTTCTCAGTGGCGACCTCCACCAGCTCCAGATCCTGCTGAGAGATGTTGAA
GGTCTTCATGATGGCCCTCCTATAGTGAGTCGTATTATACTATGCCGATATACTATGCCGATGATTAATTGTCAACT
TGGTAACGAATCAGACAATTGACGGCTTGACGGAGTAGCATAGGGTTTGCAGAATCCCTGCTTCGTCCATTTGA
CAGGCACATTATGCAAGCATTGCTGGAACACTTTATTACCCAATCCACCGTGTATTCATTGATGGCGGTGGTGTT
GGTGGCCTTTCTGGAGTCGCTGGCGCTGGTCGGTTTGATTCTACCCGGTACGGTG

**CP6-araB-gentR**

GGCGTGGTGCCCAGCGGTTTCAACACCAGTTTGCAGGGCAGTTTCGCTTCCGTATTCAGCGCATTAACGACGT
GCTCGGCATGTTGGGTGACCTGACGCAGGGTTTCCGGGCCATACAGATGCTGGCTGCCAATGACAAACCACA
CTTCATAATTATCAAAAATCGTCATTATCGTGTCCTTAGGTGGCGGTACTTGGGTCGATATCAAAGTGCATCACTT
CTTCCCGTATGCCCAACTTTGTATAGAGAGCCACTGCGGGATCGTCACCGTAATCTGCTTGCACGTAGATCACA
TAAGCACCAAGCGCGTTGGCCTCATGCTTGAGGAGATTGATGAGCGCGGTGGCAATGCCCTGCCTCCGGTGC
TCGCCGGAGACTGCGAGATCATAGATATAGATCTCACTACGCGGCTGCTCAAACTTGGGCAGAACGTAAGCCG
CGAGAGCGCCAACAACCGCTTCTTGGTCGAAGGCAGCAAGCGCGATGAATGTCTTACTACGGAGCAAGTTCC
CGAGGTAATCGGAGTCCGGCTGATGTTGGGAGTAGGTGGCTACGTCTCCGAACTCACGACCGAAAAGATCAA
GAGCAGCCCGCATGGATTTGACTTGGTCAGGGCCGAGCCTACATGTGCGAATGATGCCCATACTTGAGCCACC
TAACTTTGTTTTAGGGCGACTGCCCTGCTGCGTAACATCGTTGCTGCTGCGTAACATATGTTTTTCCTCCTTATG
TTAAGCTTACTCAGTTATTATATCATAAATATCTGTGTCAAGAATAAACTCCCACATGGATTCGGAAGTTCCTATAC
TTTCTAGAGAATAGGAACTTCCTATTATAGAGTCGCAACGGCCTGGGCAGCCTGTGCCGGGGCGGAAGTTGGA
AGATAGTGTTGTTCGGCGCTCATCGCCCATTGCTGATAGCGGCGATAAAGCTGTTCAAAGCGTTGTGCCTGCT
CGCTGCACGGTTGCAGGGTTTTCTCTACCGCACTGGCCATTTTTTGCTGAGCTGATGGGATGTCTGCGTGCAC
TTTCGCGGCGACGGCAGCAAAAATCGCCGCACCGAGCGCACAGCACTGGTCAGAGGCAACAATTTGCAGCG
GGCGATTCAGCACGTCGCAGCAGGCCTGCATAATGACCTGGTTTTTCCGCGCGATGCCGCCCAGTGCCATCA
CGTTATTAACGGCGATCCCCTGATCGGTAAAGCACTCCATGATTGCGCGTGCGCCAAAGGCGGTGGCAGCAAT
CAAACCGCCGAACAGCAGCGGAGCGTCGGTAGCGAGGTTAAGATCGGTAATCACCCCTTTCAGGCGTTGGTT
AGCGTTCGGTGTGCGGCGGCCGTTAAACCAGTCGAGCACCACCGGCAGGTGATCCAGAGACGGATTTTTGG
CCCATGCTTCGGTCAGCGCCGGAAGCAGTTGTTTCTGGCTGGCGTTGATTTGCGTTTTCAGTTCCGGATGCTG
GGCGGCAAGCTGTTCCAGCGGCCAGCCGAGTACGCGACCAAACCAGGCGTAGATATCACCAAACGCCGATTG
GCCTGCTTCCAGACCGATAAATCCAGGCACCACGCTGCCATCAACCTGACCGCAAATACCTTTAACTGCCCGC
TCGCCAACGCTCTGTTTGTCGGCAATCAGAATGTCGCAGGTGGAAGTACCGATAACTTTTACCAGTGCGTTAG
GCTGTGCGCCTGCGCCAACTGCGCCCATATGGCAGTCAAACGCGCCGCCGGAAATCACCACGCTTTCAGGCA
GGCCGAGACGCTGCGCCCATTCCGGGCATAAGGTGCCCACCGGAATATCGGCAGTCCAAGTGTCAGTGAACA
GCGGGGAAGGCAAATGGCGATTGAGGATCGGGTCCAGCTCATCAAAGAAACTGGCTGGCGGCAGGCCGCCC
CAGCTTTCGTGCCACAGAGATTTATGCCCGGCGCTGCAACGTCCGCGACGAATATCCTGCGGGCGGGTGGTA
CCGGAAAGCAGAGCTGGCACCCAGTCGCACAGCTCAATCCACGATGCGGCAGATTGCGCCACGGCGCTGTC
CTGGCGAGTCACATGCAGGATTTTTGCCCAGAACCATTCGCTGGAATAAATACCACCAATGTAGCGGGAGTAGT
CAACGTTGCCCGGCGCGTGGCACAAACGGGTAATCTCTTCCGCTTCTTCAACCGCAGTGTGGTCTTTCCACAA
TACGAACATCGCGTTCGGGTTTTCGGCAAACTCCGGGCGCAGCGCCAGCACGTTTCCGTCGGCATCAATCGG
TGCGGGCGTCGAGCCGGTACTGTCAACGCCAATCCCGACCACAGCTGCGCGCTGTTCGACGCTAAGCTCTGC
AAGCACGGTTTTCAGTGCCGCTTCCATTGACTCAATGTAGTCACGCGGATGATGACGGAACTGGTTATTCGGG
GCATCACAAAATTGCCCTTTCTGCCAACGGGGATACCACTCTACGCTGGTGGCGATCTCTTCACCGGTAGCGC
AGTCCACCGCCAAAGCTCGCACAGAATCACTGCCAAAATCGAGGCCAATTGCAATCGCCATCGTTTCACTCCA

| |
|---|
| TCCAAAAAAACGGGTGTCGTATTATACTATGCCGATATACTATGCCGATGATTAATTGTCAACTTGGTAACGAATC<br>AGACAATTGACGGCTTGACGGAGTAGCATAGGGTTTGCAGAATCCCTGCTTCGTCCATTTGACAGGCACATTAT<br>GCAAGCATTGCTGGAACACTTTATTACCCAATCCACCGTGTATTCATTGATGGCGGTGGTGTTGG |
| **araE-bsdR** |
| CTACGCTGGTGGCGATCTCTTCACCGGTAGCGCAGTCCACCGCCAAAGCTCGCACAGAATCACTGCCAAAATC<br>GAGGCCAATTGCAATCGCCATCGTTTCACTCCATCCAAAAAAACGGGTTTACTTTAGTTCCTGGTGTACTTGAG<br>GGGGATGAGTTCCTCAATGGTGGTTTTGACCAGCTTGCCATTCATCTCAATGAGCACAAAGCAGTCAGGAGCA<br>TAGTCAGAGATGAGCTCTCTGCACATGCCACAGGGGCTGACCACCCTGATGGATCTGTCCACCTCATCAGAGT<br>AGGGGTGCCTGACAGCCACAATGGTGTCAAAGTCCTTCTGCCCGTTGCTCACAGCAGACCCAATGGCAATGG<br>CTTCAGCACAGACAGTGACCCTGCCAATGTAGGCCTCAATGTGGACAGCAGAGATGATCTCCCCAGTCTTGGT<br>CCTGATGGCCGCCCCGACATGGTGCTTGTTGTCCTCATAGAGCATGGTGATCTTCTCAGTGGCGACCTCCACC<br>AGCTCCAGATCCTGCTGAGAGATGTTGAAGGTCTTCATGATGGCCCTCCTATAGTGATTACACACCGATATTGC<br>GCAGCTTCTCGCCGGCCATCAGTTTACGCTCAATATGTTCAAGGGTCACGTTCTTCGTCTCCGGGATCAGCCA<br>AAATGTAATGCCTACGAACGCAATATTCAGCGCCGTATATAACCAAAAGGTTCCCGCCGCACCAATGCTATCTAA<br>CAGGGTCAGGAACGTGGCGCCGATAATCATATTGCTAACCCAATTGGTCGTGGTTGAACAGGTGATGCCAAAG<br>TCGCGACACTTCAGTGGTTGAATTTCGCTACACAAAATCCAAACTACCGGTGCCGCCGACATGGCGTAGCCGG<br>CAATACACATCATGGTCATGCCCACCGACAACCAAGACAGTCCAGAACTAGCCGTCCCGTTATCGAACTGCATC<br>AGGCAATAGCCTAACACAAGGGTTCCCAGGGCCATAACGCTGAAGCCAATCTTCAGAGCCGGTTTACGGCCCG<br>CCTTATCTACAGTAAATACCGCAATAAAGGTCGCGAACATGAACGTAAGACCAACGACCAAGGTCGCGATCATC<br>TGCTGTTCCGTTGTAGTAAAACCCGCCATCTTAAAAATGCGCGGGGCATAATACATAATAATATTCATTCCAGTGA<br>ATTGCTGCATCGCCTGCAGCAGCATGCCCAGAAAAACGGCACGGCGGACATTGCGATTAATTTTAAACAGCGC<br>CCAGCCCCCCTGCTTCAGTTTCAGCGATTCGCGGATCTCGTTCAGCTCTTCACGCGCCTTTTCTGACGTATCG<br>CGTAACATGCGCAGCACTTCTTCCGCTTCAATATGACGGCCCTTCTCCGCCAGCCAGCGCGGAGAATTCGGCA<br>GAAAAACAACTAAAATGATCAGCAGGACCGCCGGCAGGGCTAACACGCCCAGCATCGCACGCCAATTACCTGA<br>GTAAGAAAAGGCCGTATCCGACAGGAATGCTAAAACAATACCAAGGGTGACCATAAGTTGATACATGCTGATCAT<br>CTTACCGCGTACGTTCTCGCTCGCCATTTCGGACAGGTAAAGAGGCGCGGTGTAACTCGCGATGCCGACGGC<br>GATCCCTAATACAACGCGTGCTGCGATCAGCATCTCGACCGAAGTGGCAAACGCGCTGCCGATGGAACCAAGT<br>ACGAAAAGAATCGCGCCGGCCATCAAGCTATACTTACGACCCAGGCGGAAACTCAACCAGCCGTTGAACAGG<br>GCCCCAATCGCCGCACCCAGCATCATTGAGCTAACTACCCATTCCTGCAGACGGCTGGTAAGCACAAAGTGAT<br>CCGTGATGAATGGTAACGCCCCCGCGATAACACCAATATCTAATCCAAACAGCAGCCCTGCAACGGCGGCTGC<br>AACCGATACAAACATGTTCATACGGCGCGTGTCACGTAAGCTACGTGGTGTAAGGGCGGATTCCGTGTTAATGG<br>TGACCATAATCAACCTCCTTACGACAGTATATCTGATTGTCGTATTATACTATGCCGATATACTATGCCGATGATTA<br>ATTGTCAACTTGGTAACGAATCAGACAATTGACGGCTTGACGGAGTAGCATAGGGTTTGCAGAATCCCTGCTTC<br>GTCCATTTGACAGGCACATTATGCAAGCATTGCTGGAACACTTTATTACCCAATCC |

**Table S3:** Oligos used for strain confirmations & sequencing

| **Oligo** | **Sequence** |
|---|---|
| glf_intF | GCATCAGAAGGGCTTATTGG |
| pdxK_intR | GAATTGATTTTGCCGCCGCTG |
| ampR_intF | CCGCATACACTATTCTCAGAA |

| | |
|---|---|
| araA_intF | GATAACGACACACGCCTGCC |
| polB_intR | CATTGTGCATATCACCCTCG |
| araB_intF | GCTGTTCGACGCTAAGCTC |
| yabI_intR | CAACACCACCGCCATCAATG |
| araE_intR | GATCTCGTTCAGCTCTTCACG |

**Table S4:** Composition of media used for robustness study

| **Name** | **Glucose (g $L^{-1}$)** | **Xylose (g $L^{-1}$)** | **Arabinose (g $L^{-1}$)** | **Galactose (g $L^{-1}$)** |
|---|---|---|---|---|
| G20 | 9 | 29 | 4.6 | 2.3 |
| G30 | 13.5 | 25.4 | 4.1 | 2 |
| G40 | 18 | 21.8 | 3.5 | 1.7 |
| G50 | 22.5 | 18.1 | 2.9 | 1.5 |
| G60 | 27 | 14.4 | 2.3 | 1.2 |
| G70 | 31.5 | 10.9 | 1.7 | 0.9 |
| G80 | 36 | 7.3 | 1.2 | 0.6 |
| H1 | 9 | 32.4 | 1.8 | 1.8 |
| H2 | 9 | 25.2 | 7.2 | 3.6 |
| H3 | 9 | 19.8 | 10.8 | 5.4 |

| | | | | |
|---|---|---|---|---|
| H4 | 9 | 18 | 12.6 | 5.4 |
| H5 | 9 | 16.2 | 16.2 | 3.6 |
| H6 | 9 | 14.4 | 18 | 3.6 |
| H7 | 9 | 10.8 | 19.8 | 5.4 |
| H8 | 31.5 | 12.2 | 0.7 | 0.7 |
| H9 | 31.5 | 9.5 | 2.7 | 1.4 |
| H10 | 31.5 | 7.4 | 4.1 | 2 |
| H11 | 31.5 | 6.8 | 4.7 | 2 |
| H12 | 31.5 | 6.1 | 6.1 | 1.4 |
| H13 | 31.5 | 5.4 | 6.8 | 1.4 |
| H14 | 31.5 | 4.1 | 7.4 | 2 |

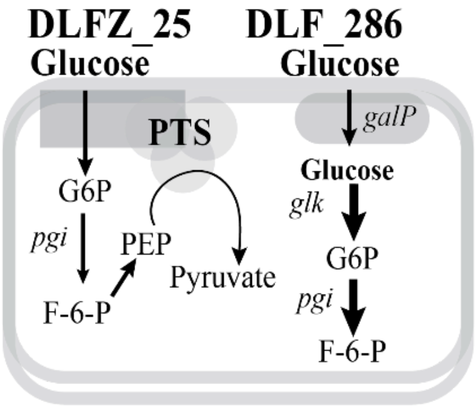


**Figure S1.** Schematic comparison of central carbon metabolism in the control strains DLF_Z0025, DLF_0286. DLF_Z0025 relies on PTS-dependent glucose uptake and glycolysis-based metabolism, whereas DLF_0286 utilizes non-PTS glucose uptake while retaining glycolysis-based central metabolism.

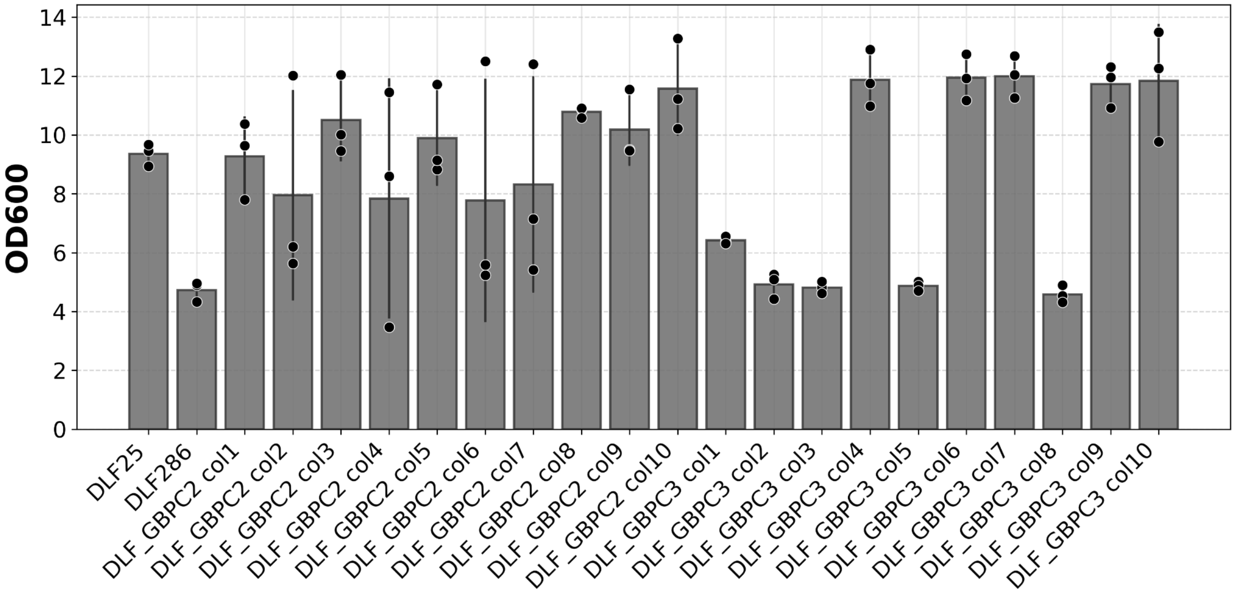


**Figure S2.** Growth profile of 10 colonies picked after construction of DLF_GBPC2 and DLF_GBPC3 strain
